# Stomata Detector: High-throughput automation of stomata counting in a population of African rice (*Oryza glaberrima*) using transfer learning

**DOI:** 10.1101/2021.12.01.469618

**Authors:** Sophie B. Cowling, Hamidreza Soltani, Sean Mayes, Erik H. Murchie

## Abstract

Stomata are dynamic structures that control the gaseous exchange of CO_2_ from the external to internal environment and water loss through transpiration. The density and morphology of stomata have important consequences in crop productivity and water use efficiency, both are integral considerations when breeding climate change resilient crops. The phenotyping of stomata is a slow manual process and provides a substantial bottleneck when characterising phenotypic and genetic variation for crop improvement. There are currently no open-source methods to automate stomatal counting. We used 380 human annotated micrographs of *O. glaberrima* and *O. sativa* at x20 and x40 objectives for testing and training. Training was completed using the transfer learning for deep neural networks method and R-CNN object detection model. At a x40 objective our method was able to accurately detect stomata (n = 540, *r* = 0.94, p<0.0001), with an overall similarity of 99% between human and automated counting methods. Our method can batch process large files of images. As proof of concept, characterised the stomatal density in a population of 155 *O. glaberrima* accessions, using 13,100 micrographs. Here, we present developed Stomata Detector; an open source, sophisticated piece of software for the plant science community that can accurately identify stomata in *Oryza spp*., and potentially other monocot species.

## Introduction

The improvement of elite crop varieties is essential in meeting increasing global food demand (Ray *et al*., 2019; FAO *et al*., 2020) and resilience to a changing climate (Kissoudis *et al*., 2016; Gao, 2021). A cornerstone in the development of crop resilience to global warming lies in balancing the trade-off between plant water relations and photosynthesis, in which stomata play a central role (Chaves *et al*., 2002; Lawson and Blatt, 2014). Stomata are dynamic microscopic pores in the leaf epidermis, which act as the gatekeepers between a leaf’s internal and the external environments. Stomata open and close in response to internal and environmental stimuli, such as light and heat. In doing so, stomata regulate the degree of CO_2_ assimilation for photosynthesis (*A*) and water lost through transpiration, both occuring during stomatal conductance (*gs*) (Lawson and Blatt, 2014; Kostaki *et al*., 2020; Yang *et al*., 2020). However, there is a trade-off between improved plant water use efficiency (WUE = *A/gs*) and CO_2_ assimilated for crop productivity (Blum, 2009; Lawson *et al*., 2010; Lawson and Blatt, 2014). Modifying the density of stomata on the leaf surface (SD) is one method that plants use to balance *A* with WUE, for example high levels of SD have been shown to enhance photosynthetic rate by 30% in *Arabidopsis thaliana* (Tanaka et al. 2013), while a reduction of SD in rice (*Oryza sativa*) showed an improvement in WUE without compromising photosynthesis and yield (Caine *et al*., 2019; Mohammed *et al*., 2019).

As a functional trait, understanding the effects of stomatal density is also important to several other plant research areas. The speed of stomatal dynamics of opening and closing places a limitation on photosynthetic efficiency under fluctuating light conditions, with consequences for both WUE and productivity and speed may be in part determined by stomata size (Drake *et al*, 2013; Lawson and Blatt, 2014). Stomata are also known to be key players in mediating pathogen resistance, where density and anatomy effect the likelihood of colonisation (Melotto *et al*., 2008; McKown *et al*., 2014; Muir, 2020). Consequently, stomatal density and morphology are an important research focus in the search for improved crop productivity and resilience in future climates.

Stomatal density has been shown to be a heritable trait, controlled by suites of genes, such as *EPIDERMAL PATTERNING FACTORs (EPFs), EPF-LIKEs* and the *ERECTA-family* (Hara *et al*., 2007; Hara *et al*., 2009; Hunt and Gray, 2009; Zoulias *et al*., 2018). The genetic control of stomatal development and patterning is relatively well-defined in *Arabidopsis thaliana* (Nadeau and Sack, 2002; Chowdhury *et al*., 2021). Elucidating the genetic control of stomatal features has only begun in crops relatively recently (McAdam et al. 2021), which is a major barrier when harnessing stomatal traits for crop improvement. Considering the important role that stomata play in carbon acquisition and water use efficiency, the successful introgression of diverse drought and heat resilient plant genotypes is essential to future crop improvement (Qu *et al*., 2016; Faralli, Matthews and Lawson, 2019; Kimura *et al*., 2020; Moore *et al*., 2021). To achieve this, large scale stomatal phenotyping efforts are necessary to capture the diversity across crop species, their wild relatives and, ideally, the wider plant kingdom. This information can then be used to identify desirable stomatal traits, elucidate the underlying genetic control and the genetic diversity through genomic analysis for future breeding programs.

The scientific resources and characterisation of genomic research has rapidly evolved, becoming increasingly precise and high throughout. Whereas rapid, large scale trait measurement pipelines are sorely lacking and recognised to be a major data collection bottleneck. This limits crop breeding progress in global food and nutritional security (Mir *et al*., 2019). The measurement of stomatal morphology and patterning is a slow, manual process, comprising of three main steps; 1. acquisition of leaf surface impressions, usually via dental putty (non-destructive) or clear nail varnish (destructive but faster); 2. image acquisition using a microscope, obtaining up to 20 fields of view per peel, based on the microscope objective, peel quality and stomatal size; 3. Measurement of stomatal density and morphology, often completed using the image analysis software ImageJ (National Institute of Health, USA). Manual processing is an acceptable technique when working on small numbers of individuals but proves a significant limitation when measuring stomatal traits in a large population, which is necessary for documenting stomatal diversity and genomic analyses. As mentioned above, genomic analyses are increasingly used for identifying trait-related candidate loci and the underlying genetic variation in a trait of interest but require large numbers of individuals for accurate loci detection (Hong and Park 2012).

The analysis of epidermal and stomatal traits is manual, slow, fraught with human error and limits whole population phenotyping efforts. And while there have been recent developments in this area (Laga, Shahinnia and Fleury, 2014; Duarte, De Carvalho and Martins, 2017; Jayakody *et al*., 2017), there are currently no established, open-source automated stomatal analysis methods available to hasten the process. Of the methods that have been developed, none can accurately identify stomata across variable taxa and do not reliably capture the core crop species (wheat, rice, maize), that will be a focus in securing future food security. There is now an urgent need to develop automated, high-throughput phenotyping of stomatal traits for the scientific community, to benefit both fundamental research and crop improvement. In this project, we aim to substantially reduce the time, cost and error associated with the manual counting of stomata through the development of an automated counting method. This project uses a machine-learning based method to accurately count stomata from a microscopy image. As large volumes of images are produced during a population level phenotyping effort, the method can process a file of multiple images, as well as single micrographs. The accuracy and useability of the method is then tested by phenotyping stomatal density in a population of *O. glaberrima* accessions, consisting of 155 genotypes. Here, we develop a resource that will improve research standards and provide information on stomatal traits at a population level and assist in the identification of molecular markers in the selection of new crop varieties for a sustainable future.

## Methods

### Plant material and growth

The seed of 155 *O. glaberrima* accessions was provided by Diversité Adaptation Developpement des plantes (DIADE), IRD-Montpellier, France (Supp. Table 1). Plants were grown and measured in a controlled environment agronomy style glasshouse (Cambridge HOK, UK) at the Sutton Bonington Campus, University of Nottingham, UK. Conditions were maintained at 28±3 °C, 50-60% relative humidity and a 12-hour dark:light (07.00 – 19.00 hrs) photoperiod. Light levels were controlled using blackout blinds and metal halide lamps were used to maintain light levels when they fell below 200 mmol m^-2^ s^-1^ photosynthetically active radiation (PAR). Seeds were heat treated to prevent pathogenesis at the primary seedling stage through immersing in water at 55°C for 15 minutes. Seedlings were grown in module trays and transplanted to soil pits (5mx5mx1.25m, LxWxD) within the glasshouse at 2 weeks old. Five replicates of each accession were transplanted in east – west rows (Supp. Fig. 1), at 20cm intervals, into high nutrient loam-based soil. Accessions were grown and planted in rotations of twelve genotypes at a time, staggered at 1–2-week intervals. Plants were measured at eight weeks old, and four accessions were selected for measurement. Measurements commenced July 2017 and finalised October 2017 (Supp. Table 2). The *O. sativa* variety ‘IR64’ was used as a reference genotype, as this is well characterised, and planted as a row in every batch.

### Stomatal impressions and image collection

Impressions of the leaf epidermis were taken from an approximately 1cm^2^ mid-section of the first fully expanded leaf, using fast drying clear nail polish and adhered to a microscope slide. Impressions of the abaxial (basal) and adaxial (upper) leaf surface was taken, as stomatal density and morphology can differ between the two leaf sides (Franks and Farquhar, 2007; Chatterjee *et al*., 2020).

Images were obtained on a Leica DM5000B light microscope at x20 and x40 objectives, using the image settings; 70% brightness, 0.35 gamma and greyscale. Images were saved as a 1728×1944 intermediate quality resolution pdf, this was a compromise between image quality and size for computational processing speed. Microscopy images of impressions for the entire experimental panel of 155 *O. glaberrima* accessions and *O. sativa* ‘IR64’ replicates were obtained just using the x40 objective, with 10 fields of view per impression. This produced a total of 13,110 micrographs to process for phenotypic analysis.

### Image annotation

Image annotation is a necessary process in training a computer vision model to accurately identify an object within an image. Our process included the manual definition, using a bounding-box, and a text-based description of a stoma within a micrograph. The generation of annotated stomata microscopy images for training was completed using the open source ‘Labellmg’ software (Tzutalin, 2015) (Fig. 1). 380 images at x20 and x40 objectives were annotated, of which 20% was used for testing and 80% for training.

**Figure 1:**
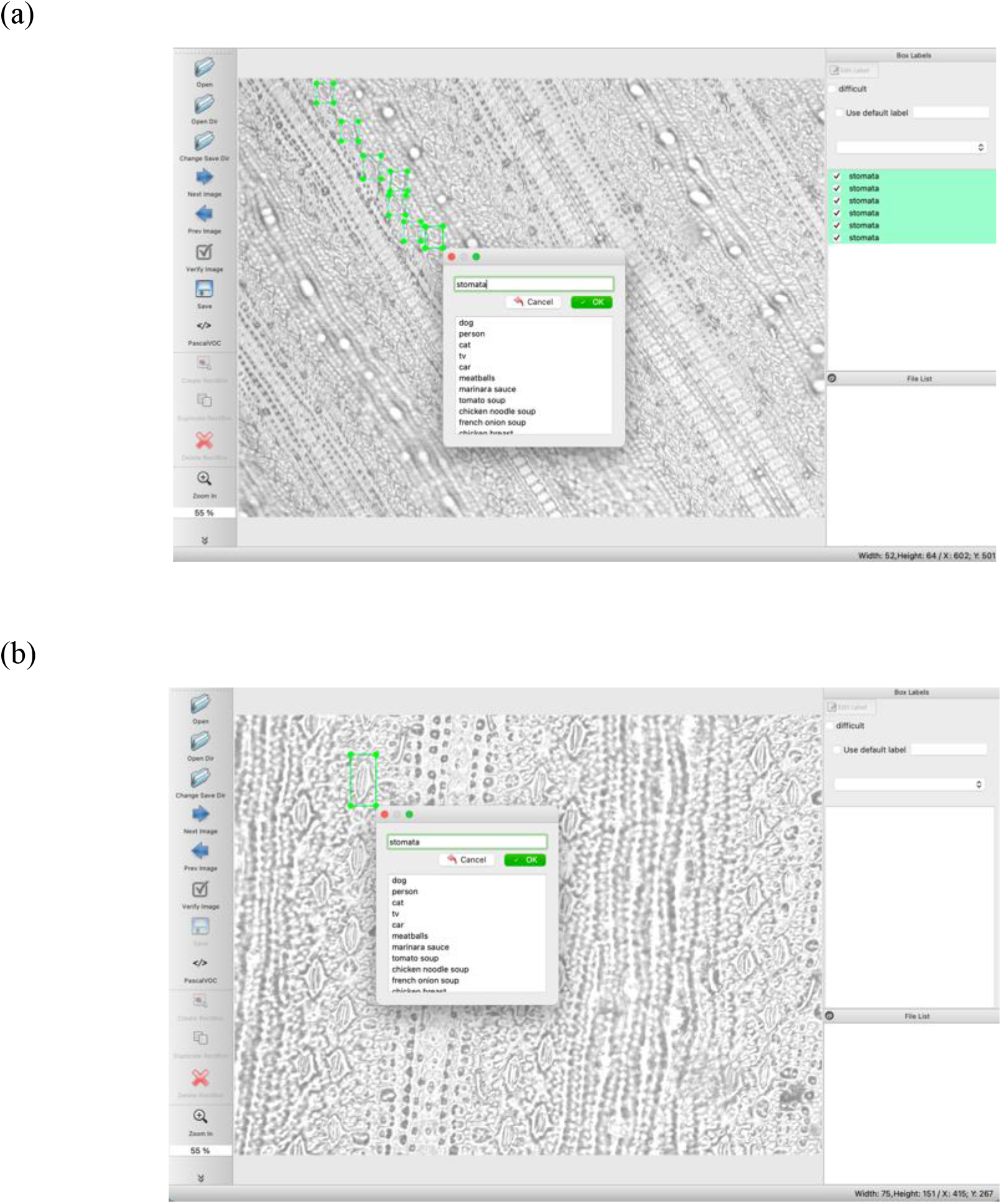
Image annotation using the graphical image annotation tool, Labellmg. Microscopy images of the rice species *O. glaberrima* and *O. sativa* were annotated at (a) x20 and (b) x40 objectives using Labellmg software.

Post software development, the accuracy of the software and experimental purpose was tested using the *O. glaberrima* population. Testing the accuracy of the software was completed on a subset of microscopy images at x20 and x40 microscope objectives. See Supp. Fig. 2 for images of the Stomata Detector software.

### Transfer learning

Our method is based on transfer learning for deep neural networks. That is, we have utilised a pre-trained deep model for the different datasets and adapt it for our stomata samples. Here, we briefly review the transfer learning technique and provide the details of the used model in our experiment.

Transfer learning is a machine learning technique in which a learner learns a new task using the experiences of another task but related to the new task. It is also a popular approach in deep learning as it helps to significantly reduce the training time and complexity of designing a deep model resulting in a well-trained model for the target task with good generalisation performance. Transfer learning is used extensively in computer vision and natural language processing. For instance, if the target task is the classification of the CIFAR-10 images, it is better to use a pre-trained model on a very large dataset such as ImageNet as a starting point and adapt this pre-trained model for the new task (e.g., classifying of CIFAR-10 images) by only training a small part of the model. In this way, not only can we use the features learned by a related task, but also, we do not need to design and start training parameters of a model from scratch. This is indeed one of the important features of modern deep learning frameworks such as TensorFlow and PyTorch which provide a rich class of pre-trained models as a built-in API. This helps practitioners and researchers to quickly train an existing model for their own setup to test the performance.

### Object Detection Model

Based on the transfer learning approach, we utilise a pre-trained object detection model trained on the standard COCO datasets (Lin et al. 2014). This dataset is a collection of more than 330k images with 80 object categories for large-scale object detection, segmentation, and captioning tasks. Since our goal is detecting and classifying stomata, we use the Faster R-CNN model (Ren *et al*., 2015) as one of the state-of-the-art methods based on deep neural networks. In particular, we downloaded a pre-trained Faster R-CNN model available in Tensorflow with the Inception-V2 architecture (Szegedy et al. 2016) as the base model. Inception-V2 is a variation of Inception-V1 also referred to as GoogLeNet was the state-of-the-art architecture at ImageNet competition in ILSRVRC 2014. After loading the pre-trained Faster R-CNN, the last few layers of classification layers are changed to meet the aim of stomata classification and detection. In the next step, the Faster R-CNN with stomata images are trained with different hyper parameters such as learning rate and number of epochs to find out the best parameters to reduce execution time and errors.

All statistical analysis was completed in R-Studio (v. 4.0.5), graphs were generated using the R package ggplot2 (v. 3.3.3).

### Stomata Detector software functionality

A user-friendly interface was created (see Supp. Fig. 2 for screenshots) to facilitate automated stomatal detection for end users that are not literate in handling and running code. Stomata detector requires a threshold value to be set before analysing stomatal micrographs. This is the value the requires an inputted value of 0-1, this is the estimated certainty that the algorithm has correctly identified an object in the micrograph as a stoma (1 = 100% certainty). The user can then decide to load and execute a singular or file of many micrographs. The main interface shows a list of loaded micrographs on the right-hand panel, a summary of the historical and current analyses in the central window. A window is also generated, showing the code in the backend of the software, this can be useful when monitoring the progress when batch processing a file of micrographs. When an analysis is complete, a .csv output file is generated with columns detailing; the micrograph file path name, the chosen threshold value, number of detected stomata, number of stomata not included if they did not pass the defined threshold value, mean and standard deviation.

## Results

### Stomata Detector can rapidly batch process micrographs

The Stomata Detector software can batch process a folder and singular micrographs. When a single micrograph is analysed by the software, the output includes both a .csv file of results and an image of the original micrograph overlaid with bounding boxes over objects that are identified as stomata (Fig. 2c - d and Supp. Fig. c). When sequentially batch processing a file of images, a .csv file of the results is still produced but to save on processing time and storage capacity, the image overlaid with bounding boxes is not. The software can batch process high numbers of images in a comparably short period of time, for example a folder of all 13,110 *O. glaberrima* stomatal impression micrographs was processed within 24 hours. Single images, with accompanied image outputs, are processed within microseconds.

**Figure 2:**
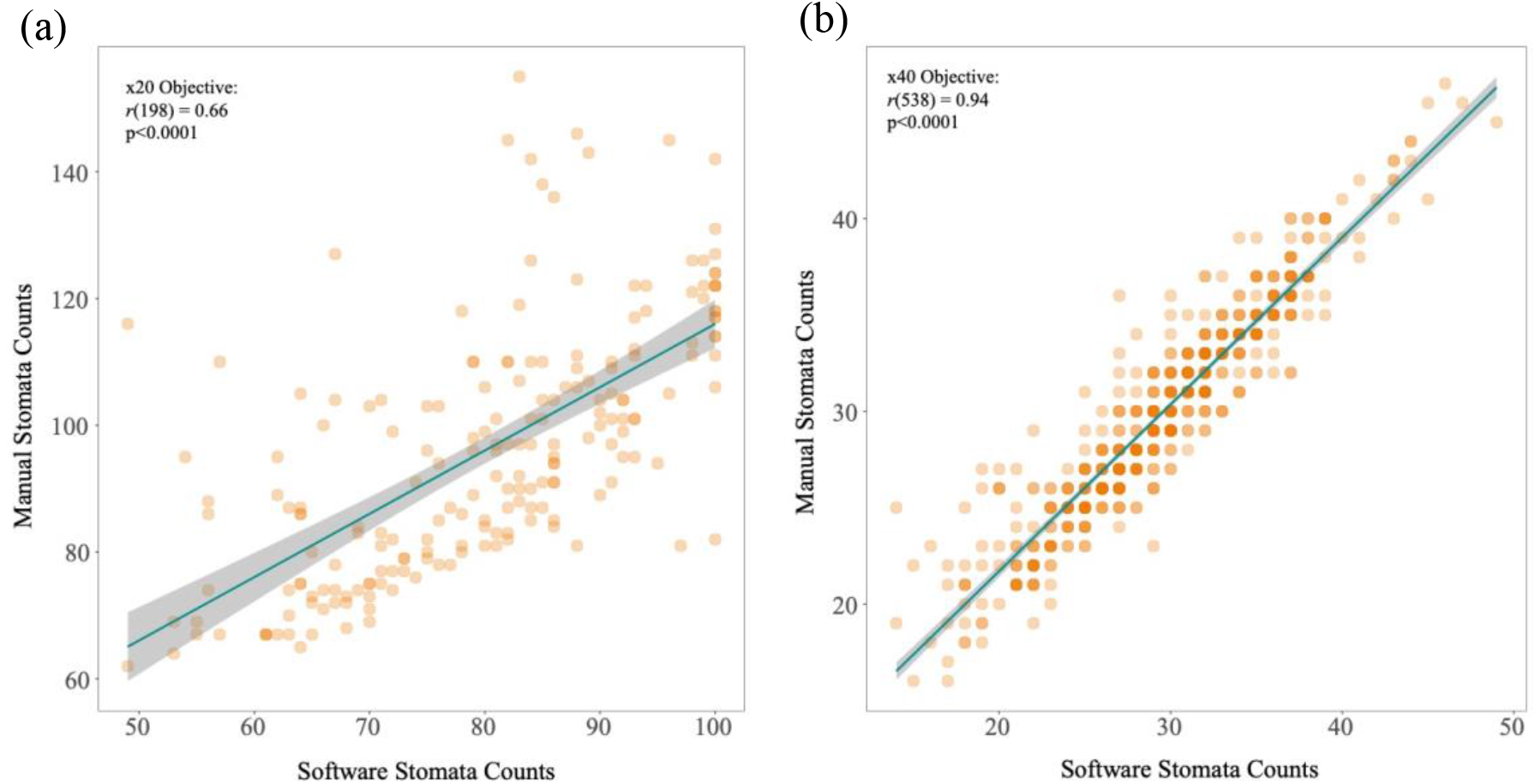
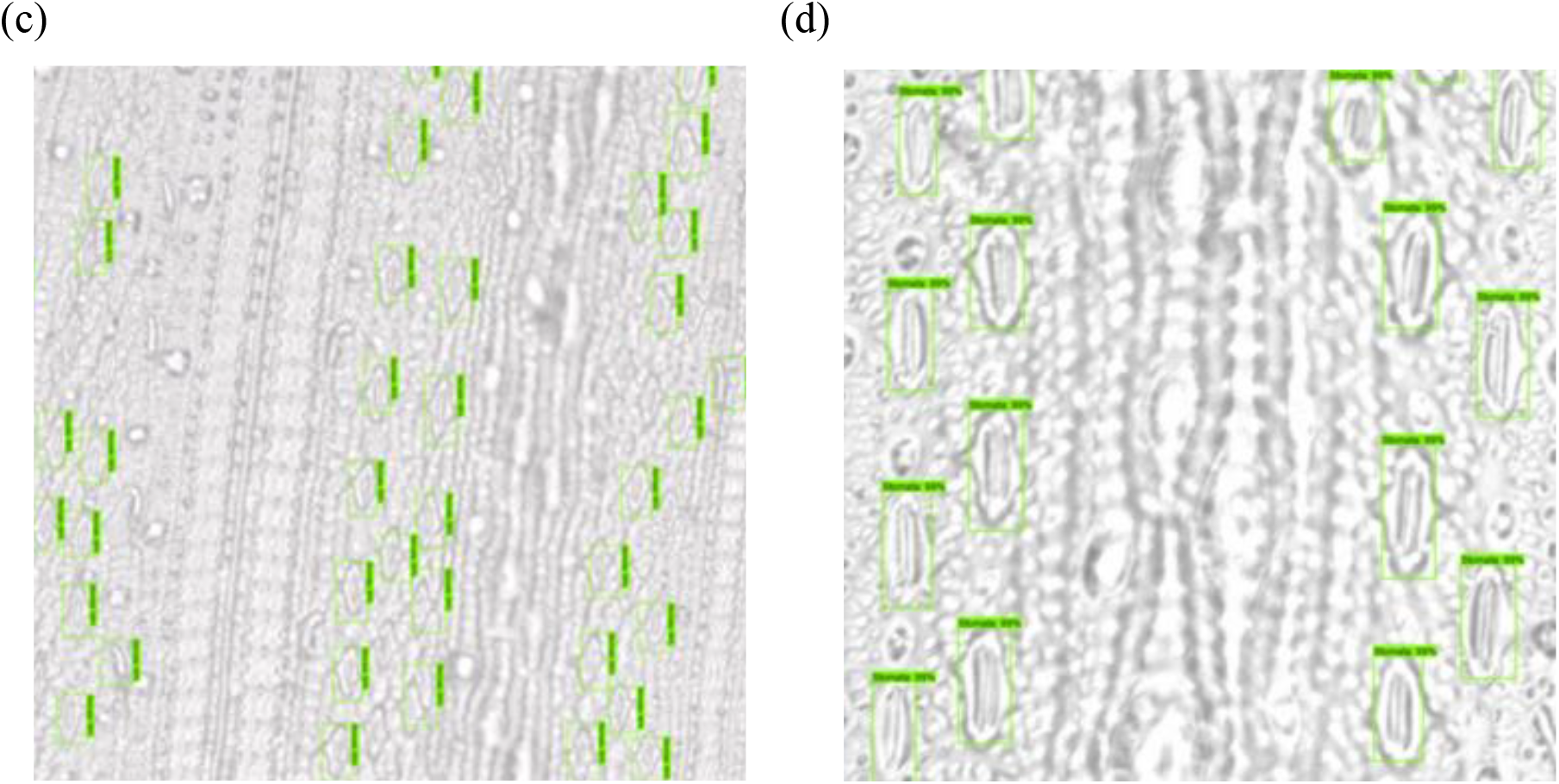
Stomata Detector identifies stomata better at higher magnifications. The stomata count obtained from the automated software method was compared to the manual counting method for x20 (a; *r* = 0.66, p<0.0001) and (b; *r* = 0.94, p<0.0001) x40 microscope objectives. Images (c) and (d) show cropped Stomata Detector outputs, detected stomata are shown bound in green boxes.

### Stomatal detection accuracy

The software was tested for accuracy on images of rice (*O. glaberrima* and *O. sativa*) stomata at x20 and x40 objective, to establish the accuracy at each magnification. The correlation between the automated Stomata Detector and manual counting methods were used as one indication of software accuracy. At x20 magnification a Pearson correlation test showed a significant association between manual and automated stomatal counts (n = 200, *r* = 0.66, p<0.0001; Fig. 2a and c), with an overall similarity of 83% between the two stomatal counting methods. While a higher correlation was observed at a x40 objective (n = 540, *r* = 0.94, p<0.0001; Fig. 2b and d), with an overall similarity of 99% between the total sum of two stomatal counting methods. A total of 20 micrographs, 10 for each leaf side, were randomly selected from the pool of 13,100 *O. glaberrima* population micrographs. These were used to further check the accuracy of the automated analysis method in comparison to manual stomata counting (Fig. 3a-b and Table 1). Counts matched for 12/20 micrographs between the two methods, with an overall similarity of 99.8% for the total stomatal counts (*r* = 0.94, p<0.0001). The remainder 8/20 micrographs showed a small difference in stomata count between manual and automated stomatal counting methods (Table 1). Of these, stomata were missed in 3/20 and false positives identified in 5/20 micrographs. Examples of false positive and negative stomata identification from these images can be seen in Fig. 4 (a-b).

**Figure 3:**
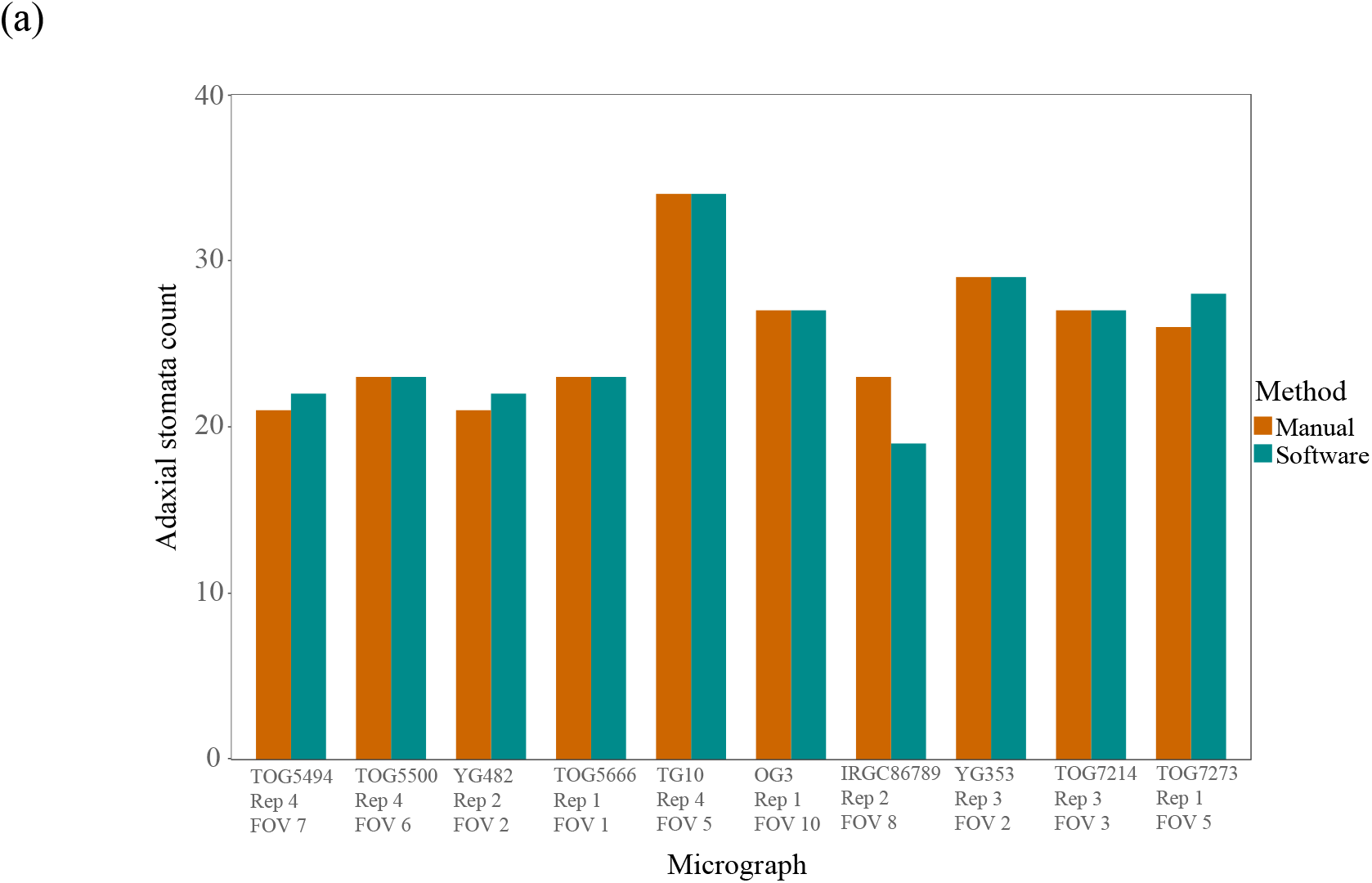
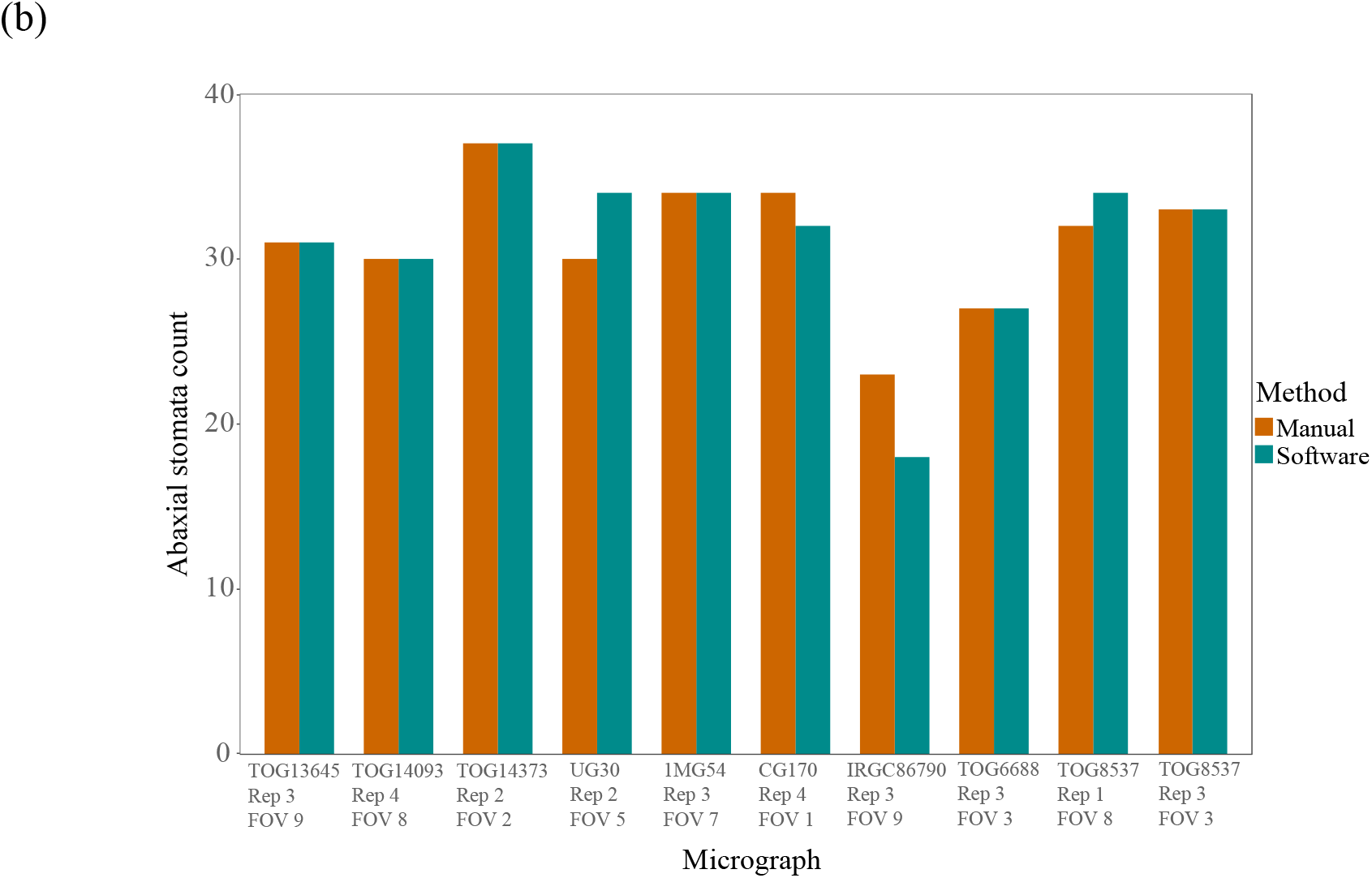
Bar charts showing the comparison of manual and automated software stomata counting methods from randomly selected micrographs. 20 outputs, taken from the analysis on 13,100 micrographs from a population of 155 *O. glaberrima* accessions, using the automated stomata counting method were randomly selected ((a) 10 adaxial, (b) 10 adaxial). The micrographs of the randomly selected outputs were manually counted to compare the accuracy between the two methods.

**Figure 4:**
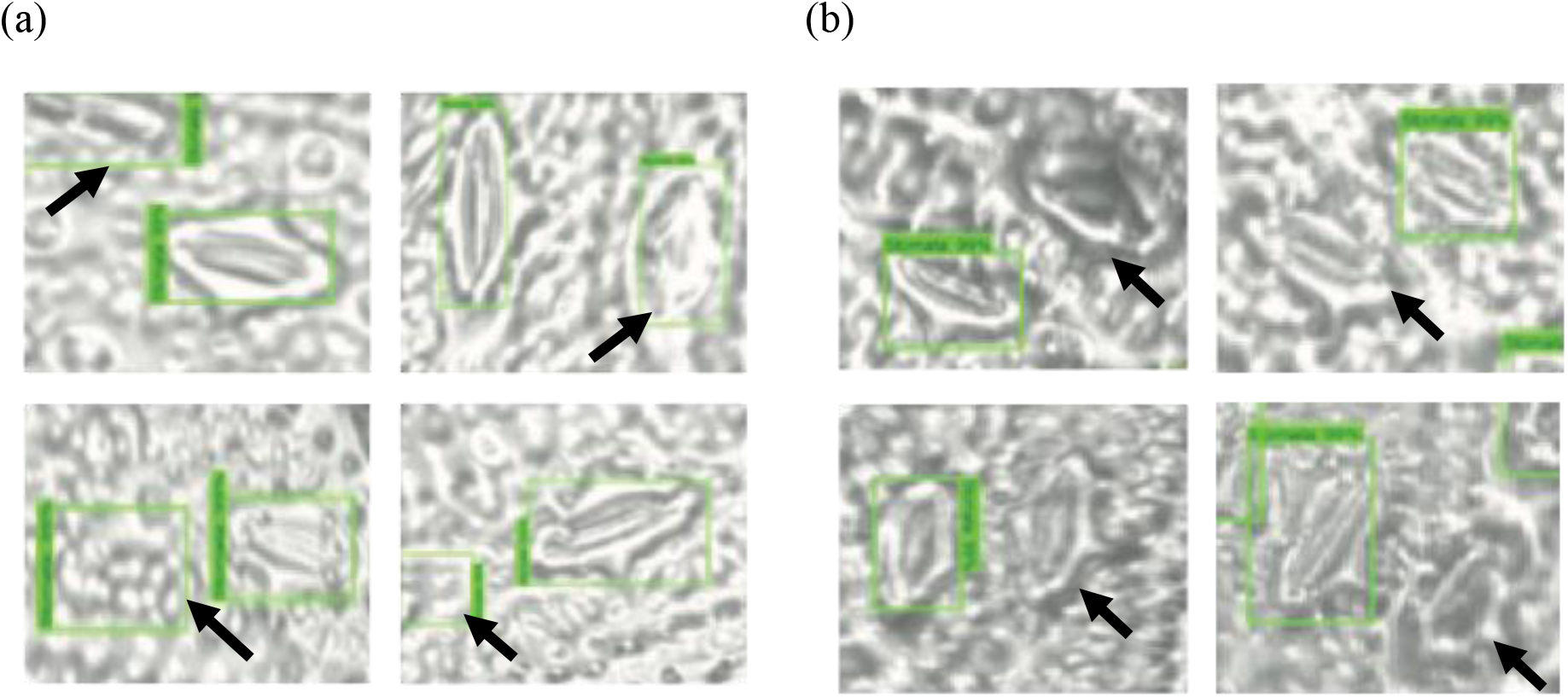
Samples showing errors in stomatal detection. False positive (a) identification occurs when features in the micrograph are erroneously detected as stomata. While false negatives (b) are stomata detected by a human during manual counting but not by the automated method. Black arrows indicate a false positive and false negative occurrence.

**Table 1:**
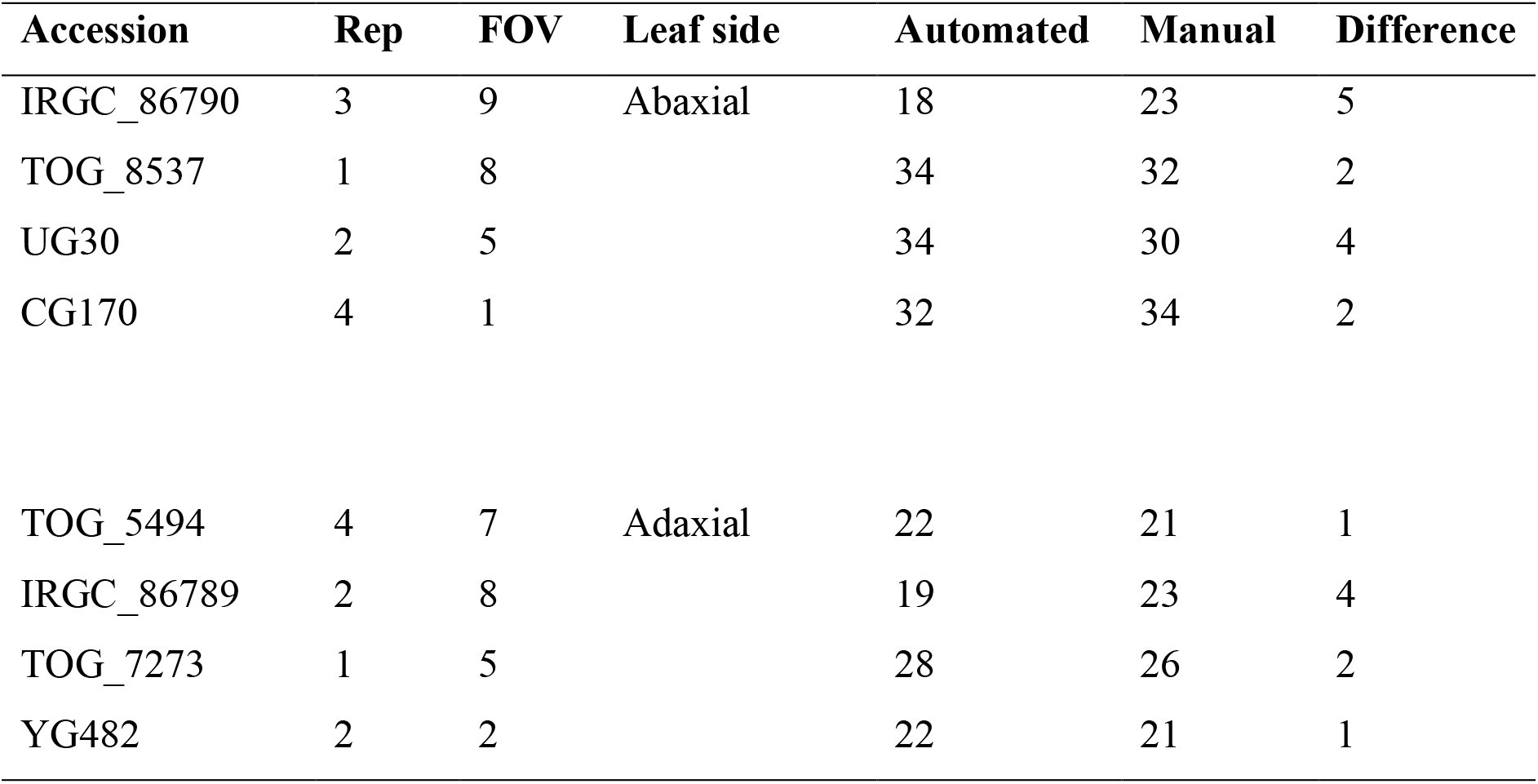
Randomly selected micrographs showing a difference in stomata count between automated and manual methods. The table shows the micrograph identifying name (accession name, biological replicate (rep) and field of view (FOV)), the stomata count for the stomata counting method and the difference between these two methods.

### Quantifying variation in a population of 155 individuals

As proof of concept, we used Stomata Detector to process 13,100 micrographs, collected from a population of 155 *O. glaberrima* accessions and the *O. sativa* cultivar, IR64. This population was previously uncharacterised for stomatal traits. While the micrographs were obtained by hand, the automated analysis was completed within 24 hours. The stomatal density (per mm^2^) was calculated and showed a normal trait distribution, with no spurious outliers (Fig. 4). The stomatal density values for the population (abaxial = 377 mm^-2^, adaxial = 305 mm^-2^) are in line with that reported by Chatterjee *et al*. (2020) for *O. glaberrima* accession IRGC_103544. Supp. Fig. 3 shows boxplots of the stomatal density for each accession, calculated from the outputs of Stomata Detector.

**Figure 4:**
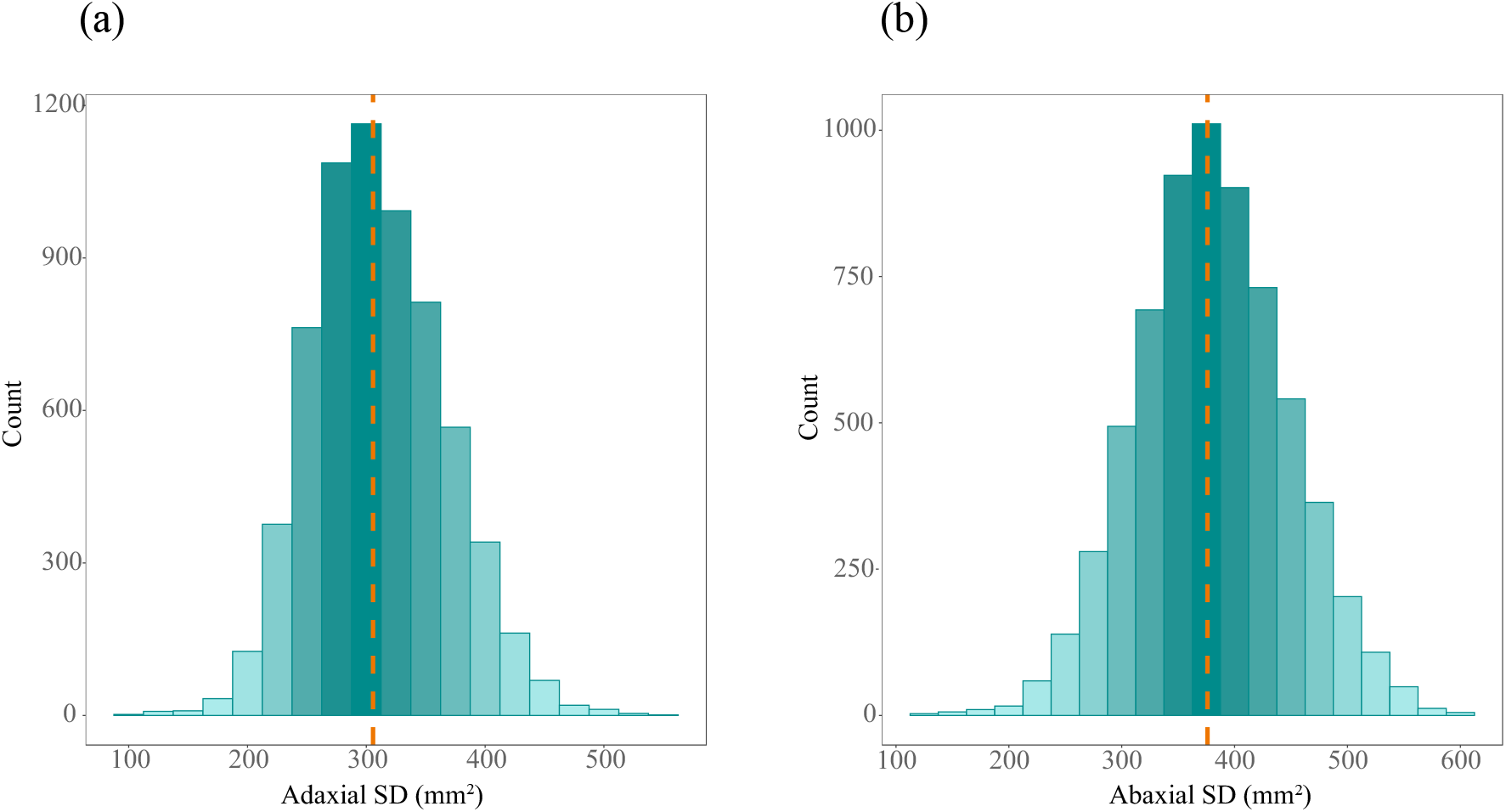
Trait distribution of (a) adaxial and (b) abaxial stomatal density for the *O. glaberrima* population, analysed by the automated method. Orange line shows the mean value in the trait distribution.

## Discussion

We have a better understanding than ever that stomatal density and morphology is a critical functional trait for optimising carbon assimilation and minimising abiotic stress (Caine *et al*., 2019; Faralli, Matthews and Lawson, 2019). As commercial crop species are genetically narrow (Meyer and Purugganan 2013) and poorly adapted to challenging environmental conditions, the focus is upon underutilised crop species and wild relatives as a source of stomatal genetic diversity (Meyer and Purugganan 2013). However, manual stomatal phenotyping is slow and thus limits the characterisation of stomatal genetic diversity within populations. The automatic machine-learning based software (Stomata Detector) developed here analyses the stomatal number from micrographs, that would otherwise not be feasible when manually phenotyping a large population.

A prime example of this is the *O. glaberrima* population described here, which was collected as a resource for *O. sativa* crop improvement (Cubry et al. 2020). However, the stomatal diversity in this population remains uncharacterised, which limits the elucidation of the underlying trait genetic diversity. Using the automated counting method developed here, we have been able to characterise the stomatal density in a population of *O. glaberrima* accessions. This will have a direct impact when exploring the role of stomata on *O. glaberrima* physiology, environmental resilience and elucidate trait-related gene loci.

We have shown the Stomata Detector method can accurately identify stomata, though errors can occur. False positive stomatal identification may happen when there is a structure in the micrograph that is a similar size and shape to a stoma (Fig. 3a). Observed false positives included air bubbles and the contours of an epithelial cell, particularly when on the edge of a micrograph. False negatives, that being stomata not detected by the method, typically occur when images are blurry, have low contrast between epithelial and stoma subsidiary cells or very small stomata that can be present on leaf veins. When testing this method, we compromised on image quality in favour of high throughput processing, though this may have caused occurrences of false positive and negatives. Ensuring high quality sample preparation and obtaining micrographs with a greater pixel resolution will minimise these errors, though there will be an increase in processing time. Despite this guidance, perfect stomatal peels are not always possible, and the end user requires a method that is able to accurately identify stomata in a range of peel and micrograph qualities. This can be improved by increasing the sample size, and variability of image quality, used in the training data set. The importance of choosing the optimal magnification for stomatal identification was apparent in Fig. 2a-b. This shows that Stomata Detector performed accurately at x40 objective and comparatively poorly at x20, relative to human manual analysis. It is worth noting, the comparison of this automated method to human manual counting (Fig. 2a-b) does not account for human error, which may be substantial when analysing hundreds of micrographs and is difficult to quantify. For example, the 20 randomly selected micrographs (Fig. 3; Table 1) was manually checked three times to ensure accuracy and reduce human error. 5 out of 20 images stomata were mis-counted on the first repetition of manual analysis. Stomata Detector was more accurate at a higher magnification due to the enhanced clarity and size of the leaf epidermis and stomata. Stomata are particularly small in *Oryza spp*., in comparison to other species. At the x20 objective, the leaf epidermal surface area covered is greater but means that the varnish peel across that area undulates more than at a higher magnification, causing many stomata to be out of focus and unidentifiable. Therefore, it is advised that end users dedicate thought and time to establishing the best microscope magnification, image size and quality for the species and peel quality at hand.

The Stomata Detector training set was currently enriched with images of *O. glaberrima* and *O. sativa* and does not yet reliably identify other species. There is currently no other resource that accurately identifies *Oryza spp*. stomata, other methods (Fetter et al. 2019) focuses on angiosperm taxa, with large dumbbell shaped stomata. We feel the method demonstrated here is still an asset to the scientific community, especially considering the importance of rice crop improvement as a staple food for billions of people. Though our aim would be to develop a resource to that enables taxonomically diverse stomatal ID. This would require the ongoing retraining of the method from future users.

Our initial work has developed an image processing software which allows accurate stomatal counting of 1000s of stomatal impression in a few days, rather than several months. We would like to extend this to a mobile application, which could be used in field in conjunction with a mobile phone microscope (Orth et al. 2018), enabling non-destructive real-time measurements. This would significantly alleviate the current stomata phenotyping bottleneck and alter the possibilities in trait characterisation, even extending to in field analysis of maximal stomatal conductance (Brown and Escombe, 1900; Dow, Bergmann and Berry, 2014). This would contribute to a major step forward in stomatal, and even wider leaf-level phenotyping.

If we are to tackle food insecurity and future sustainability, the integration of differing scientific techniques and fields is essential. Our project developed an open source, sophisticated piece of software that can accurately identify stomata in *Oryza spp*., and potentially other monocot, species. Demonstrating that deep machine learning is the ideal partner for narrowing the bottleneck of high-quality plant phenomics and consequently enabling crop improvement.

## Notes

### Competing Interest Statement

The authors have declared no competing interest.

