## Supplemental Material for "Stomata Detector: High-throughput automation of stomata counting in a population of African rice (*Oryza glaberrima*) using transfer learning"

**List of Supplementary material:**

Supplementary Table 1: List of *O. glaberrima* accession codes, country and African region of origin and ecology.

Supplementary Table 2: Table detailing sowing and measurement dates for phenotyping of *O. glaberrima* population and *O. sativa*, IR64.

Supplementary Figure 1 (a – c): *O. glaberrima* phenotyping experimental plan.

Supplementary Figure 2 (a – c): Images of Stomata Detector software.

Supplementary Figure 3 (a – b): Boxplot of adaxial and adaxial stomatal density for the *O. glaberrima* population.

**Supplementary table 1: List of *O. glaberrima* accession codes, ecology, country and African region of origin.** O. glaberrima germplasm was provided by Diversité Adaptation Developpement des plantes (DIADE), IRD-Montpellier, France. Information on accession ecology and country of origin was provided by AfricaRice.

| Accession code | Ecology | Country of origin | African region |
|---|---|---|---|
| IRGC_96726 | Irrigated lowland | Nigeria | West coast |
| TOG_5314 | Irrigated lowland | Nigeria | West coast |
| TOG_5321 | Rainfed lowland | Nigeria | West coast |
| TOG_5326 | Shallow forest swamp | Nigeria | West coast |
| IRGC_96740 | Irrigated lowland | Nigeria | West coast |
| TOG_5418 | Lowland | Nigeria | West coast |
| TOG_5424 | Rainfed lowland | Nigeria | West coast |
| TOG_5453 | Shallow forest swamp | Nigeria | West coast |
| TOG_5486 | Rainfed lowland | Nigeria | West coast |
| TOG_5494 | Rainfed lowland | Nigeria | West coast |
| TOG_5500 | Rainfed lowland | Nigeria | West coast |
| TOG_5556 | Rainfed lowland | Nigeria | West coast |
| IRGC_86764 | Irrigated lowland | Ghana | West coast |
| TOG_5666 | Rainfed lowland | Nigeria | West coast |
| TOG_5672 | Rainfed lowland | Nigeria | West coast |
| IRGC_96790 | Irrigated lowland | Nigeria | West coast |
| TOG_5681 | Rainfed lowland | Nigeria | West coast |
| TOG_5814 | Rainfed lowland | Liberia | West coast |
| IRGC_56785 | Irrigated lowland | Liberia | West coast |
| TOG_5882 | Rainfed lowland | Nigeria | West coast |
| IRGC_112568 | Irrigated lowland | Liberia | West coast |
| IRGC_86789 | Irrigated lowland | Liberia | West coast |
| IRGC_86790 | Irrigated lowland | Liberia | West coast |
| IRGC_86791 | Irrigated lowland | Liberia | West coast |
| TOG_5953 | Rainfed lowland | Nigeria | West coast |
| TOG_5969 | Irrigated lowland | Nigeria | West coast |

| | | | |
|---|---|---|---|
| TOG_6205 | Irrigated lowland | Guinea | West coast |
| TOG_6206 | Irrigated lowland | Zimbabwe | South inland |
| TOG_6207 | Irrigated lowland | Zimbabwe | South inland |
| TOG_6211 | Irrigated lowland | Nigeria | West coast |
| TOG_6220 | Irrigated lowland | Burkina Faso | West inland |
| TOG_6356 | Rainfed lowland | Liberia | West coast |
| TOG_6603 | Rainfed lowland | Liberia | West coast |
| TOG_6688 | Rainfed lowland | Liberia | West coast |
| TOG_6698 | Rainfed lowland | Liberia | West coast |
| TOG_6943 | Irrigated lowland | Sierra Leone | West coast |
| TOG_6951 | Irrigated lowland | Sierra Leone | West coast |
| TOG_7020 | Irrigated lowland | Sierra Leone | West coast |
| TOG_7047 | Irrigated lowland | Sierra Leone | West coast |
| TOG_7106 | Irrigated lowland | Mali | West inland |
| TOG_7108 | Irrigated lowland | Mali | West inland |
| TOG_5286 | Rainfed lowland | Nigeria | West coast |
| TOG_5400 | Lowland | Nigeria | West coast |
| TOG_5439 | Rainfed lowland | Nigeria | West coast |
| LG33 | Lowland | Mali | West inland |
| TOG_5464 | Rainfed lowland | Nigeria | West coast |
| TOG_5533 | Lowland | Nigeria | West coast |
| TOG_5566 | Rainfed lowland | Nigeria | West coast |
| TOG_5591 | Rainfed lowland | Ghana | West coast |
| TOG_5639 | Rainfed lowland | Nigeria | West coast |
| CG10 | Irrigated lowland | Senegal | West coast |
| TOG_7132 | Irrigated lowland | Senegal | West coast |
| TOG_7134 | Irrigated lowland | Senegal | West coast |
| TOG_5747 | Rainfed lowland | Liberia | West coast |
| TOG_5775 | Rainfed lowland | Liberia | West coast |
| TOG_5997 | Upland | Nigeria | West coast |
| TOG_7420 | Rainfed lowland | Sierra Leone | West coast |
| IRGC_103544 | Irrigated lowland | Mali | West inland |
| RAM 131 | Floating Rice | Mali | West inland |

| | | | |
|---|---|---|---|
| RAM 137 | Floating Rice | Mali | West inland |
| RAM 24 | Floating Rice | Guinea | West coast |
| RAM 48 | Floating Rice | Mali | West inland |
| RAM 55 | Floating Rice | Mali | West inland |
| RAM 77 | Floating Rice | Mali | West inland |
| CG14 | Irrigated lowland | Senegal | West coast |
| IG38 | Rainfed lowland | Côte d'Ivoire | West coast |
| TOG_14367 | Irrigated lowland | Guinea | West coast |
| YG353 | Rainfed lowland | Guinea | West coast |
| MG04 | Lowland | Mali | West inland |
| TOG_7214 | Irrigated lowland | Upland | West inland |
| CG171 | Irrigated lowland | Senegal | West coast |
| TOG_7219 | Irrigated lowland | Mali | West inland |
| IRGC_103549 | Irrigated lowland | Mali | West inland |
| TOG_10434 | Irrigated lowland | Côte d'Ivoire | West coast |
| TOG_7255 | Irrigated lowland | Chad | North inland |
| TOG_12086 | Rainfed lowland | Nigeria | West coast |
| TOG_12160 | Rainfed lowland | Nigeria | West coast |
| TOG_12188 | Rainfed lowland | Nigeria | West coast |
| TOG_12249 | Rainfed lowland | Nigeria | West coast |
| TOG_7273 | Irrigated lowland | Cameroon | West coast |
| TOG_7274 | Shallow Forest Swamp | Cameroon | West coast |
| IRGC_104589 | Irrigated lowland | Burkina Faso | West inland |
| IRGC_86826 | Irrigated lowland | Ghana | West coast |
| TOG_7406 | Rainfed lowland | Ghana | West coast |
| TOG_7451 | Irrigated lowland | Burkina Faso | West inland |
| TOG_7455 | Irrigated lowland | Burkina Faso | West inland |
| TOG_7456 | Irrigated lowland | Burkina Faso | West inland |
| TOG_7455 | Irrigated lowland | Côte d'Ivoire | West coast |
| TOG_7993 | Irrigated lowland | Nigeria | West coast |
| TOG_8049 | Irrigated lowland | Nigeria | West coast |
| TOG_8527 | Irrigated lowland | Gambia | West coast |

| | | | |
|---|---|---|---|
| TOG_8537 | Irrigated lowland | Gambia | West coast |
| TOG_8545 | Irrigated lowland | Gambia | West coast |
| TOG_9524 | Irrigated lowland | Côte d'Ivoire | West coast |
| TOG_12358 | Upland | Côte d'Ivoire | West coast |
| TOG_12366 | Rainfed lowland | Guinea-Bissau | West coast |
| TOG_12372 | Rainfed lowland | Guinea-Bissau | West coast |
| TOG_12387 | Rainfed lowland | Tanzania | East coast |
| TOG_12388 | Rainfed lowland | Cameroon | West coast |
| TOG_12399 | Upland | Guinea | West coast |
| TOG_12401 | Upland | Guinea | West coast |
| TOG_12411 | Upland | Guinea | West coast |
| TOG_12414 | Upland | Guinea | West coast |
| YG330 | Rainfed lowland | Guinea | West coast |
| TOG_13645 | Irrigated lowland | Guinea | West coast |
| TOG_13708 | Irrigated lowland | Guinea | West coast |
| TOG_14093 | Irrigated lowland | Guinea | West coast |
| TOG_14116 | Rainfed lowland | Liberia | West coast |
| TOG_14184 | Rainfed lowland | Zimbabwe | South inland |
| YG482 | Rainfed lowland | Guinea | West coast |
| TOG_14361 | Rainfed lowland | Guinea | West coast |
| TOG_14373 | Irrigated lowland | Guinea | West coast |
| TOG_14606 | Irrigated lowland | Guinea | West coast |
| TOG_14610 | Irrigated lowland | Guinea | West coast |
| TOG_7190 | Irrigated lowland | Côte d'Ivoire | West coast |
| IG05 | Rainfed lowland | Côte d'Ivoire | West coast |
| IG09 | Rainfed lowland | Côte d'Ivoire | West coast |
| IG14 | Rainfed lowland | Côte d'Ivoire | West coast |
| IG15 | Rainfed lowland | Côte d'Ivoire | West coast |
| IG16 | Rainfed lowland | Côte d'Ivoire | West coast |
| IG19 | Rainfed lowland | Côte d'Ivoire | West coast |
| IG21 | Rainfed lowland | Côte d'Ivoire | West coast |
| IG23 | Rainfed lowland | Côte d'Ivoire | West coast |
| IG35 | Rainfed lowland | Côte d'Ivoire | West coast |

| | | | |
|---|---|---|---|
| IG36 | Upland | Côte d'Ivoire | West coast |
| IG43 | Rainfed lowland | Côte d'Ivoire | West coast |
| IG47 | Rainfed lowland | Côte d'Ivoire | West coast |
| IG324 | Rainfed lowland | Côte d'Ivoire | West coast |
| EG55 | Lowland | Tanzania | East coast |
| EG85 | Lowland | Tanzania | East coast |
| UG14 | Lowland | Cameroon | West coast |
| UG20 | Lowland | Cameroon | West coast |
| UG26 | Lowland | Cameroon | West coast |
| UG28 | Rainfed lowland | Cameroon | West coast |
| UG30 | Rainfed lowland | Cameroon | West coast |
| LG07_S | Lowland | Mali | West inland |
| LG64 | Lowland | Mali | West inland |
| MG53 | Lowland | Mali | West inland |
| 1MG54 | Lowland | Mali | West inland |
| TG10 | Irrigated lowland | Chad | North inland |
| TG19_G | Irrigated lowland | Chad | North inland |
| TG25 | Irrigated lowland | Chad | North inland |
| TG57 | Irrigated lowland | Chad | North inland |
| CG45 | Irrigated lowland | Senegal | West coast |
| CG46 | Irrigated lowland | Senegal | West coast |
| CG70 | Irrigated lowland | Senegal | West coast |
| CG150 | Irrigated lowland | Senegal | West coast |
| CG156 | Irrigated lowland | Senegal | West coast |
| CG164 | Irrigated lowland | Senegal | West coast |
| CG170 | Irrigated lowland | Senegal | West coast |
| OG1 | lowland | Senegal | West coast |
| OG3 | lowland | Senegal | West coast |
| OG15 | lowland | Senegal | West coast |
| YG307 | Upland | Guinea | West coast |
| YG316 | Rainfed lowland | Guinea | West coast |

**Supplementary Table 2: Table detailing sowing and measurement dates of one hundred and fifty-five *O. glaberrima* accessions.** This table details how the *O. glaberrima* accessions were grouped into batches of twelve, each accession name was assigned a Nottingham line number for ease. The sowing date, date of the week that measurements commenced and date of the week that measurements ceased for each batch of accessions.

| ***O. glaberrima* line no.** | **Sowing date** | **Measurement start date** | **Measurement end date** |
|---|---|---|---|
| 1 - 12 | 26/04/2017 | 12/06/2017 | 16/06/2017 |
| 13-24 | 08/05/2017 | 26/06/2017 | 30/06/2017 |
| 25-36 | 22/05/2017 | 10/07/2017 | 14/07/2017 |
| 37-48 | 05/06/2017 | 24/07/2017 | 28/07/2017 |
| 49-60 | 19/06/2017 | 07/08/2017 | 11/08/2017 |
| 61-72 | 03/07/2017 | 21/08/2017 | 25/08/2017 |
| 73-84 | 17/07/2017 | 04/09/2017 | 08/09/2017 |
| 121-132 | 24/07/2017 | 11/09/2017 | 14/09/2017 |
| 85-69 | 31/07/2017 | 18/09/2017 | 22/09/2017 |
| 133-144 | 07/08/2017 | 25/09/2017 | 28/09/2017 |
| 97-108 | 14/08/2014 | 02/10/2017 | 06/10/2017 |
| 145-156 | 21/08/2017 | 09/10/2017 | 12/10/2017 |
| 109-120 | 28/08/2017 | 16/10/2017 | 20/10/2017 |

**Supplementary Figure 1:** ***O. glaberrima*** **phenotyping experimental plan.** (a) The left-hand side of the agronomy style glass house was divided into six small beds, three of these were used for the first six batches of measurements, following these all six beds were used until all accessions had been measured. (b) In each small bed twelve accessions and IR64, with five replicates, were grown. The accessions were surrounded by a border of spare seedlings, to minimise for edge effects. Accessions were not randomised to minimise plant damage and mismeasurement during LiCor analysis. Through this process, (c) a batch rotation through the available beds facilitated the growth and analysis of all accessions.

(a)

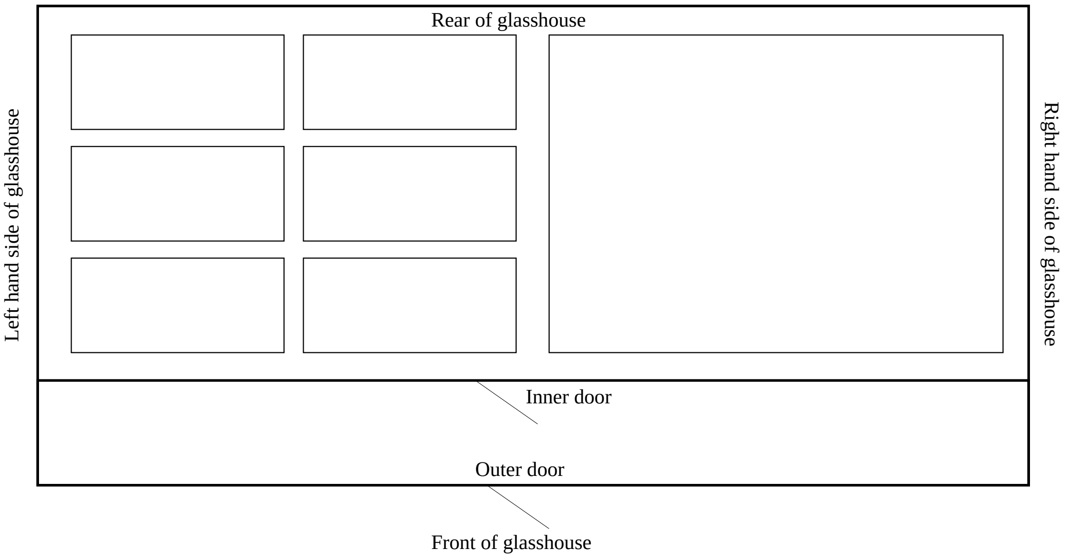


(b)

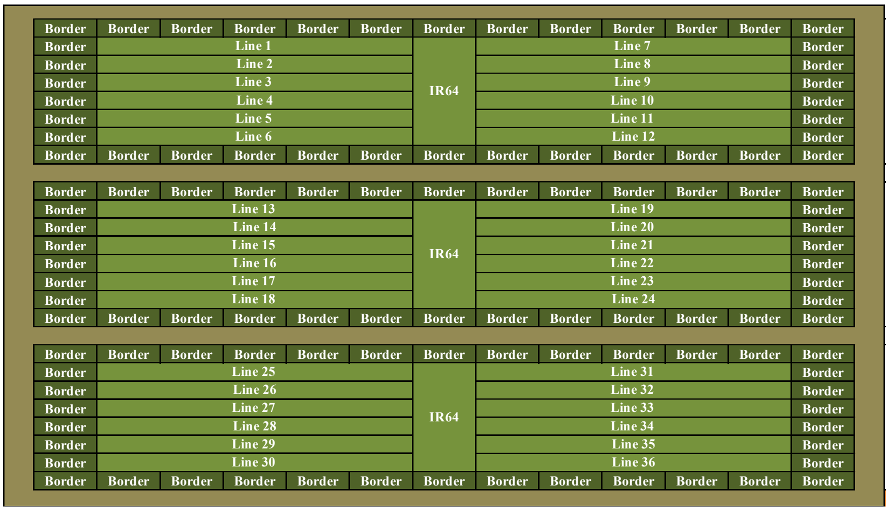


(c)

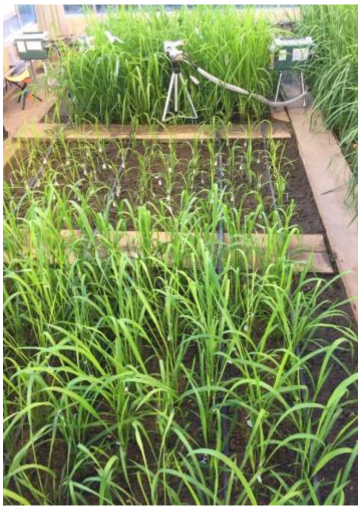

**Supplementary Figure 2: Images of Stomata Detector software.** Screenshots taken of the Stomata Detector software, showing (a) the graphical user interface, where files and folders are loaded and executed; (b) the back end, which gives a summary of which micrographs are running and the outputs; (c) image outputs from a single micrograph analysis, showing the objects identified as stomata in green bounding boxes, the total number of stomata identified and the micrograph file pathway.

(a)

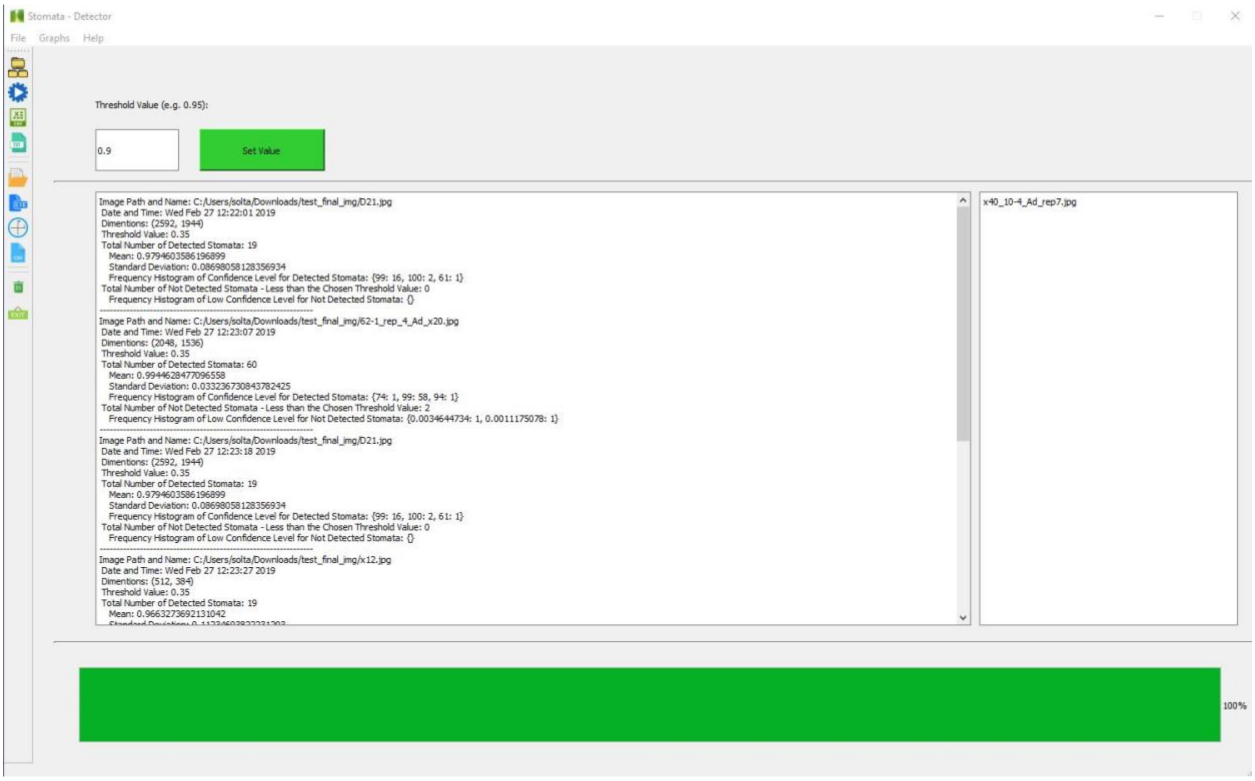


(b)

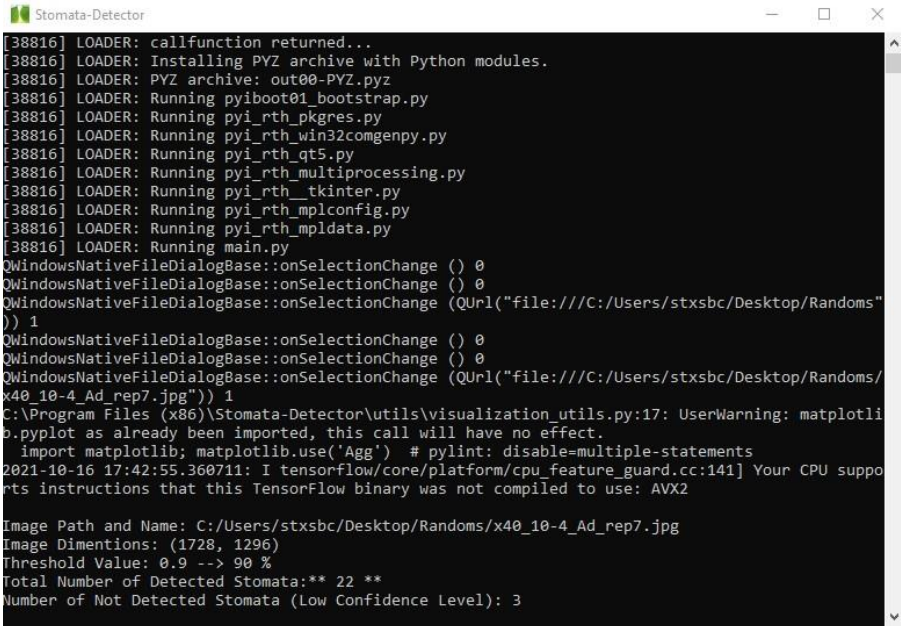

(c)

Total number of detected Stomata: 23

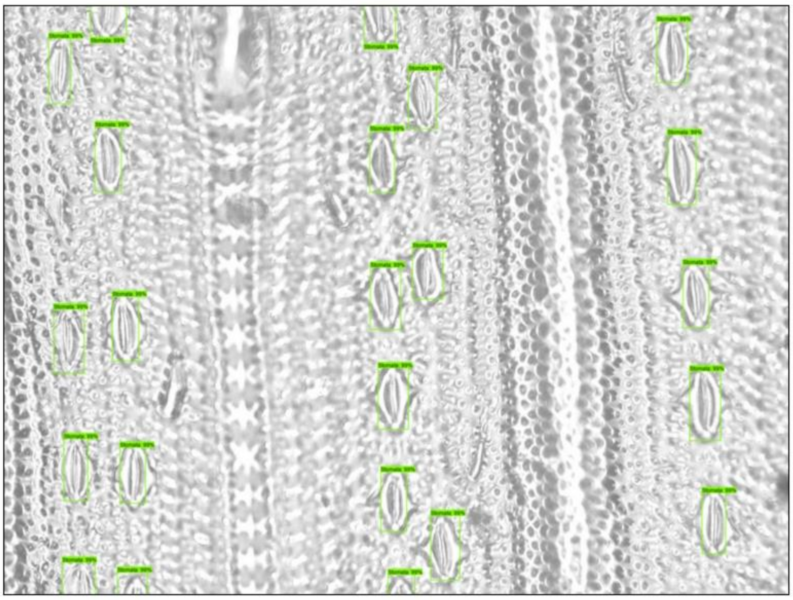

Stomata Path and Name: C:/Users/stxsbc/Desktop/Randoms/x40_11-4_Ad_rep6.jpg
Threshold value: 0.9

**Supplementary Figure 3: Boxplot of stomatal density for the *O. glaberrima* population.** The boxplots show stomatal density calculated from the automated stomata counts generated by Stomata Detector, showing (a) abaxial and (b) adaxial stomatal density for the 155 *O. glaberrima* accessions and *O. sativa* cultivar, IR64

(a)

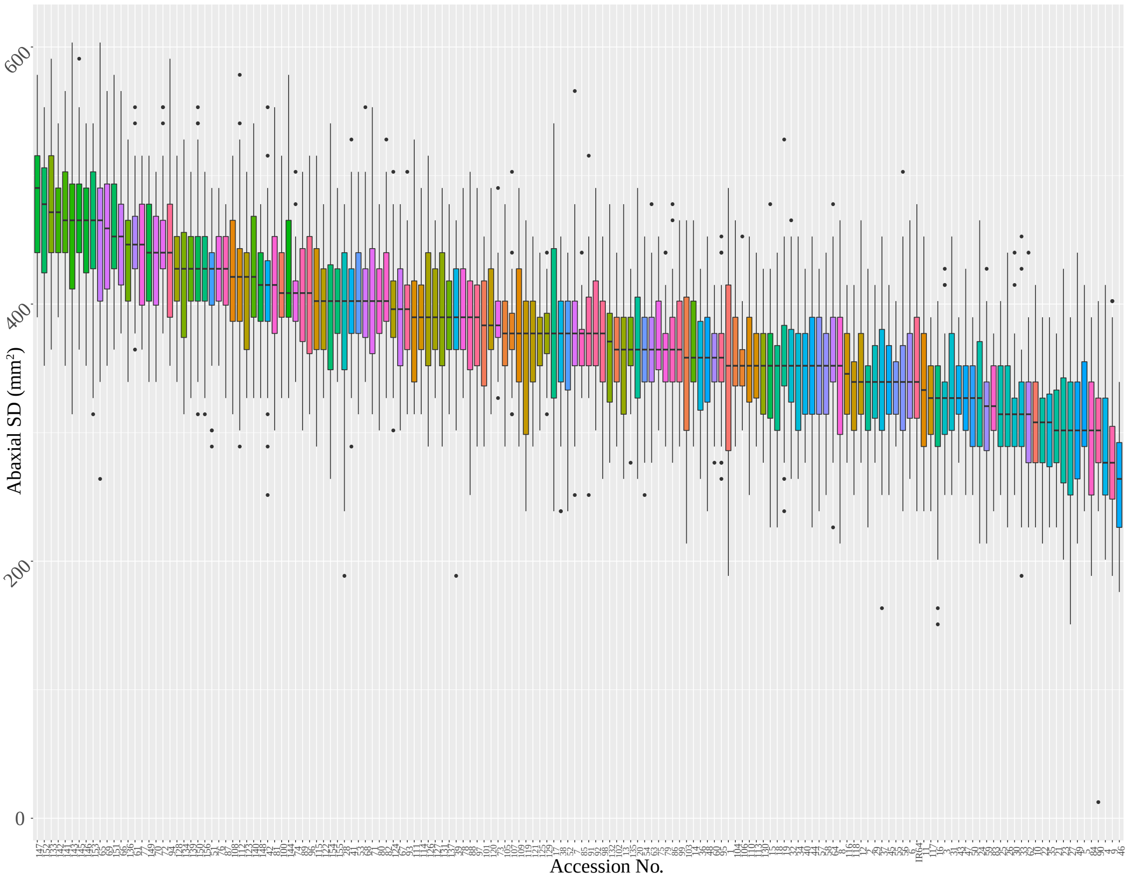


(b)

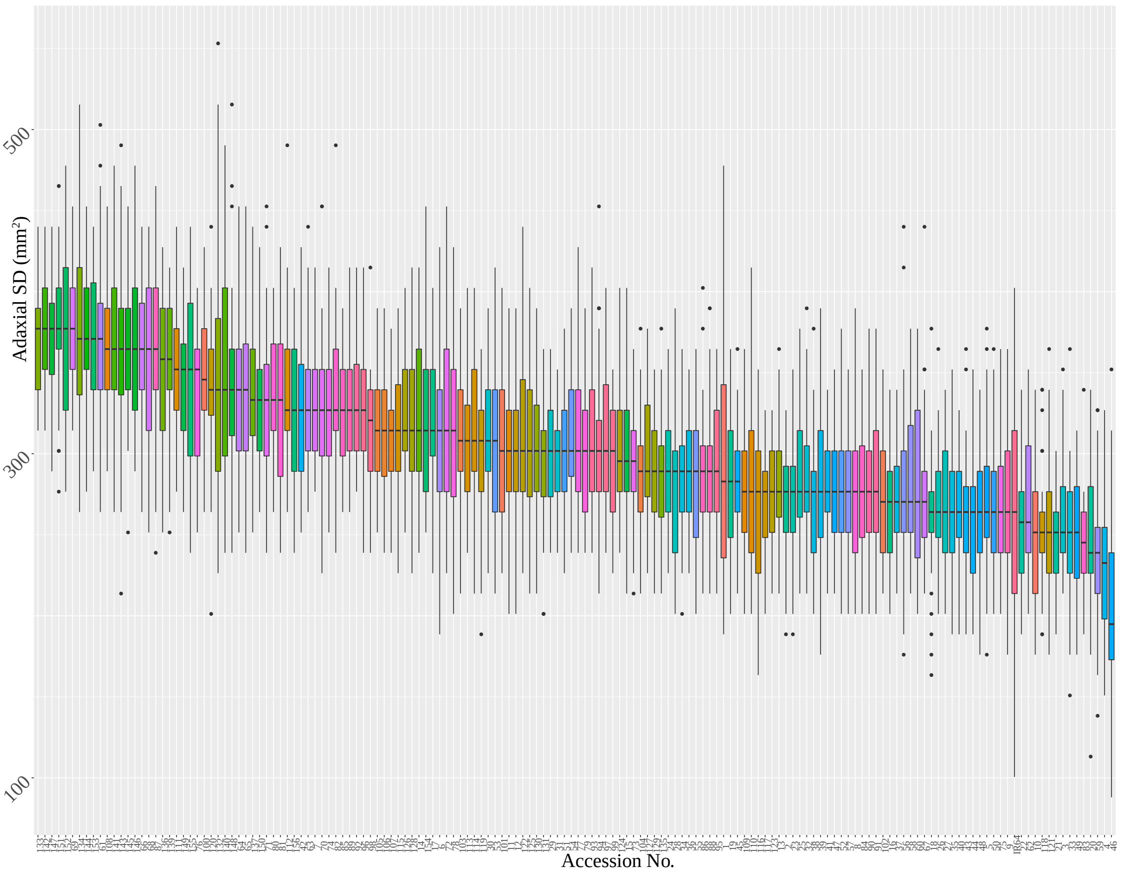
Adaxial SD (mm²)
500
300
100
Accession No.
